# Advanced Surface Passivation for High-Sensitivity Studies of Biomolecular Condensates

**DOI:** 10.1101/2024.02.12.580000

**Authors:** Run-Wen Yao, Michael K. Rosen

## Abstract

Biomolecular condensates are cellular compartments that concentrate biomolecules without an encapsulating membrane. In recent years, significant advances have been made in the understanding of condensates through biochemical reconstitution and microscopic detection of these structures. Quantitative visualization and biochemical assays of biomolecular condensates rely on surface passivation to minimize background and artifacts due to condensate adhesion. However, the challenge of undesired interactions between condensates and glass surfaces, which can alter material properties and impair observational accuracy, remains a critical hurdle. Here, we introduce an efficient, generically applicable, and simple passivation method employing self-assembly of the surfactant Pluronic F127 (PF127). The method greatly reduces nonspecific binding across a range of condensates systems for both phase-separated droplets and biomolecules in dilute phase. Additionally, by integrating PF127 passivation with the Biotin-NeutrAvidin system, we achieve controlled multi-point attachment of condensates to surfaces. This not only preserves condensate properties but also facilitates long-time FRAP imaging and high-precision single-molecule analyses. Using this method, we have explored the dynamics of polySIM molecules within polySUMO/polySIM condensates at the single-molecule level. Our observations suggest a potential heterogeneity in the distribution of available polySIM-binding sites within the condensates.

**Significance Statement:** The understanding of biomolecular condensates has significantly benefited from biochemical reconstitution with microscopy detection. Here, we present a novel surface passivation method utilizing self-assembly of Pluronic F127 on hydrophobic surfaces. This approach not only effectively minimizes non-specific binding without altering the physical properties of the condensates but also offers universal passivation across a variety of condensate systems. It demonstrates high resistance to different treatments and enables condensate immobilization through controlled anchor points. This allows for highly sensitive analytical techniques, including single-molecule imaging. The simplicity and high-performance of this method, coupled with time and cost efficiencies, could facilitate robustness and throughput of experiments, and could broaden the accessibility of biochemical phase separation studies to a wider scientific community.

## Introduction

Biomolecular condensates are cellular compartments that concentrate biomolecules without an encapsulating membrane (1, 2). Many condensates form through liquid-liquid phase separation (LLPS) of multivalent macromolecules (1, 3). Significant advancements in our understanding of condensates have derived from biochemical reconstitution with fluorescence microscopy detection (4-7). Undesirable interactions between condensates and glass surfaces represent a significant methodological challenge in these experiments. Such interactions can alter condensate material properties and generate appreciable background (4, 8, 9). Yet complete inhibition of surface adhesion is also undesirable, since many experiments are impaired by condensate movement. Effective surface passivation, balancing these effects, is crucial to robust, quantitative biochemical studies of condensates.

The most common passivation technique used in condensate biochemistry involves coating glass surfaces with methoxy polyethylene glycol (mPEG) and Bovine Serum Albumin (BSA) (4, 8). However, this method has substantial limitations. In our experience, mPEG/BSA passivation is often insufficient to prevent condensate spreading in macromolecular systems with high surface adhesion. Residual adhesion of biomolecules also produces high background that complicates high precision studies such as single-molecule imaging (10). Additionally, mPEG passivation involves many steps and is time-consuming (10-13), diminishing robustness and reproducibility, especially in laboratories without experience in condensate biochemistry. Finally surface effects typically vary between condensates and conditions, necessitating optimization when parameters change. A straightforward, efficient and generically effective passivation method is a pressing need.

Self-assembly is a spontaneous process where molecules organize into stable aggregates, driven by noncovalent interactions (14). The use of surfactant self-assembly has gained widespread recognition for its ability to inhibit non-specific biomolecule binding (15, 16). Here, we evaluated the passivation capabilities of several non-ionic surfactants on hydrophobic glass surfaces, identifying Pluronic F127 (PF127) as the most effective agent in preventing non-specific binding across both phase-separated condensates and biomolecules in dilute phases. The method exhibited excellent resilience across a broad spectrum of pH and salt conditions, while preserving the physical integrity of condensates. Combined with the Biotin-NeutrAvidin system, the method enabled condensates to be immobilized through controlled multi-point attachment to surfaces, facilitating movement-sensitive imaging, including Fluorescence Recovery After Photobleaching (FRAP) and single-molecule tracking. Using these methods, we explored the dynamics of polySIM molecules within polySUMO/polySIM condensates at the single-molecule level. Our observations suggest that the available polySIM-binding sites could have a heterogeneous distribution within the condensates.

## Results

### Screening self-assembled surfactants for glass passivation

To explore the potential of self-assembled surfactants for glass surface passivation in studies of biomolecular condensates, we tested seven non-ionic surfactants self-assembled on glass slides rendered hydrophobic by prior treatment with chlorinated organopolysiloxane (Sigmacote) (Fig. 1A). Brij L23, PF127 and Tween 20 exhibited good passivation capabilities as evidenced by formation of phase separated droplets with high contact angles on glass surfaces in experiments involving condensates of the actin regulatory proteins, Nck/N-WASP (6), and the engineered multi-valent proteins, polySUMO/polySIM (17) (Fig. 1B and 1C). These three surfactants are characterized by a hydrophobic tail, which interacts with the hydrophobic glass surface, and a polyethylene oxide-like head that forms a dense brush layer, effectively repelling molecular binding (18, 19) (Fig. 1D). Notably, PF127 exhibited the best passivation activity (Fig. 1B and 1C). Pluronic surfactants are series of commercially available triblock copolymers composed of a central polypropylene oxide (PPO) moiety flanked by polyethylene oxide (PEO) tails with various length. They stably attach to hydrophobic surfaces via adsorption of the PPO group and self-assembly of the PEO tails into a brush-like array, creating a hydrated layer that prevents close contact of other molecules with the glass surface (18). We further explored PF127 passivation in biochemical studies of condensates.

**Figure 1.**
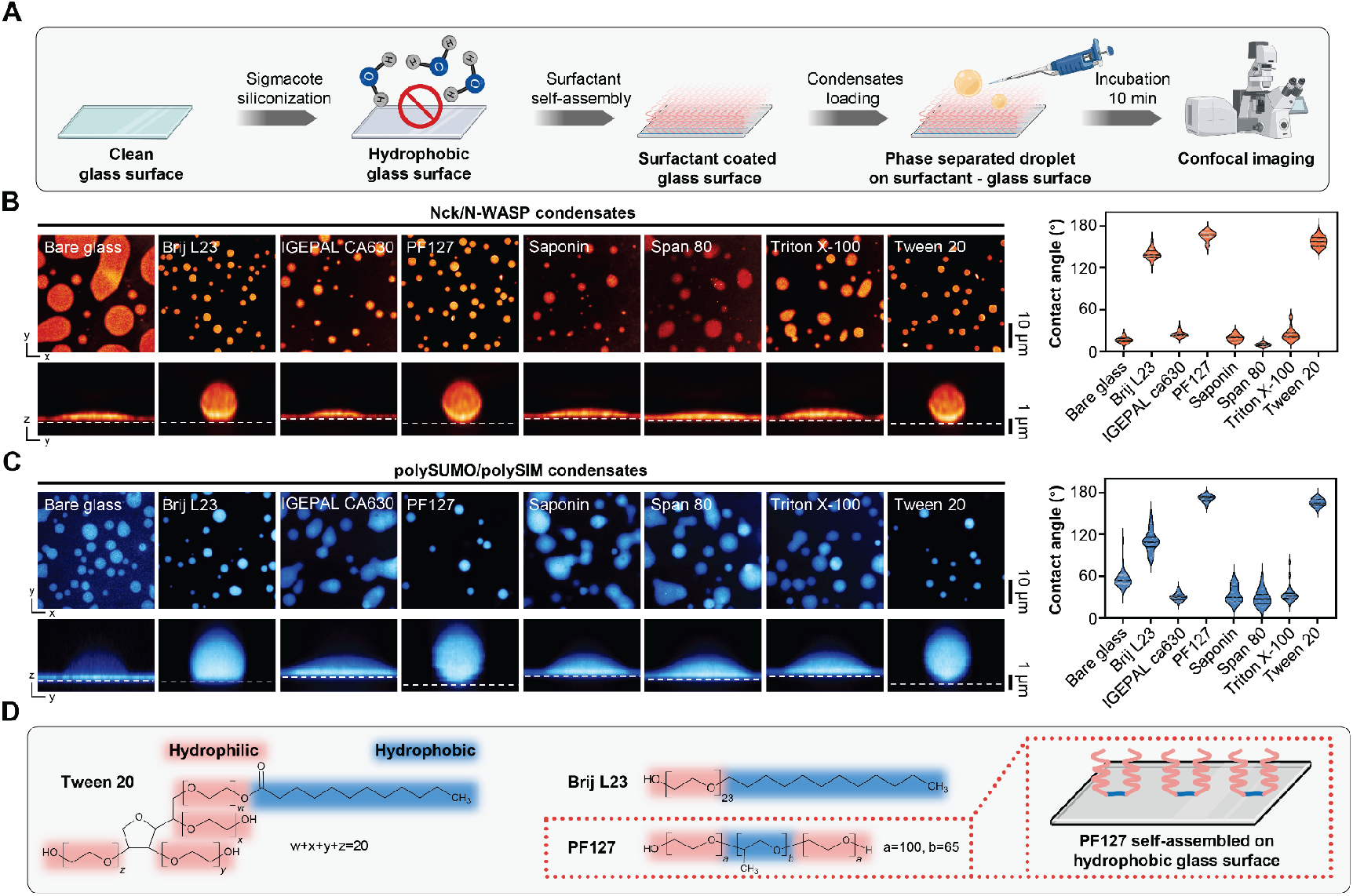
Screening self-assembled surfactants for glass passivation in phase separation studies. A, Workflow illustrating the screening process for self-assembled surfactants used in glass passivation. Image created with BioRender.com. B, Comparison of Nck/N-WASP condensates on different passivated surfaces. Plots show the contact angle (median and quartiles, n>15 condensates for each technique). C, Comparison of polySUMO/polySIM condensates on different passivated surfaces. Plots show the contact angle (median and quartiles, n>15 condensates for each technique). D, Chemical formula of surfactants demonstrating effective passivation abilities.

### PF127 passivation provides effective passivation across diverse condensate systems

The PF127 passivation procedure we developed is straightforward (Fig. 2A, Detailed Protocol in SI Appendix). The entire procedure can be completed within three hours, with active handling time <one hour (compared to >15 hours for standard mPEG/BSA treatment) (SI Appendix, Fig. S1). We compared PF127 treatment with traditional passivation techniques, including BSA, mPEG, and combined mPEG/BSA, across three condensate systems composed of: Dhh1 (7), representing IDR-mediated phase separation; Nck/N-WASP (6), generated by multivalent modular domain interactions and characterized by high wettability; and polySUMO/polySIM (17), a distinct modular domain system with low wettability. PF127 passivation yielded the lowest adhesion (highest contact angles) for all three condensates (Fig. 2B and 2C). In contrast, traditional methods produced different behaviors of each condensate on differently passivated surfaces (Fig. 2B and 2C).

**Figure 2.**
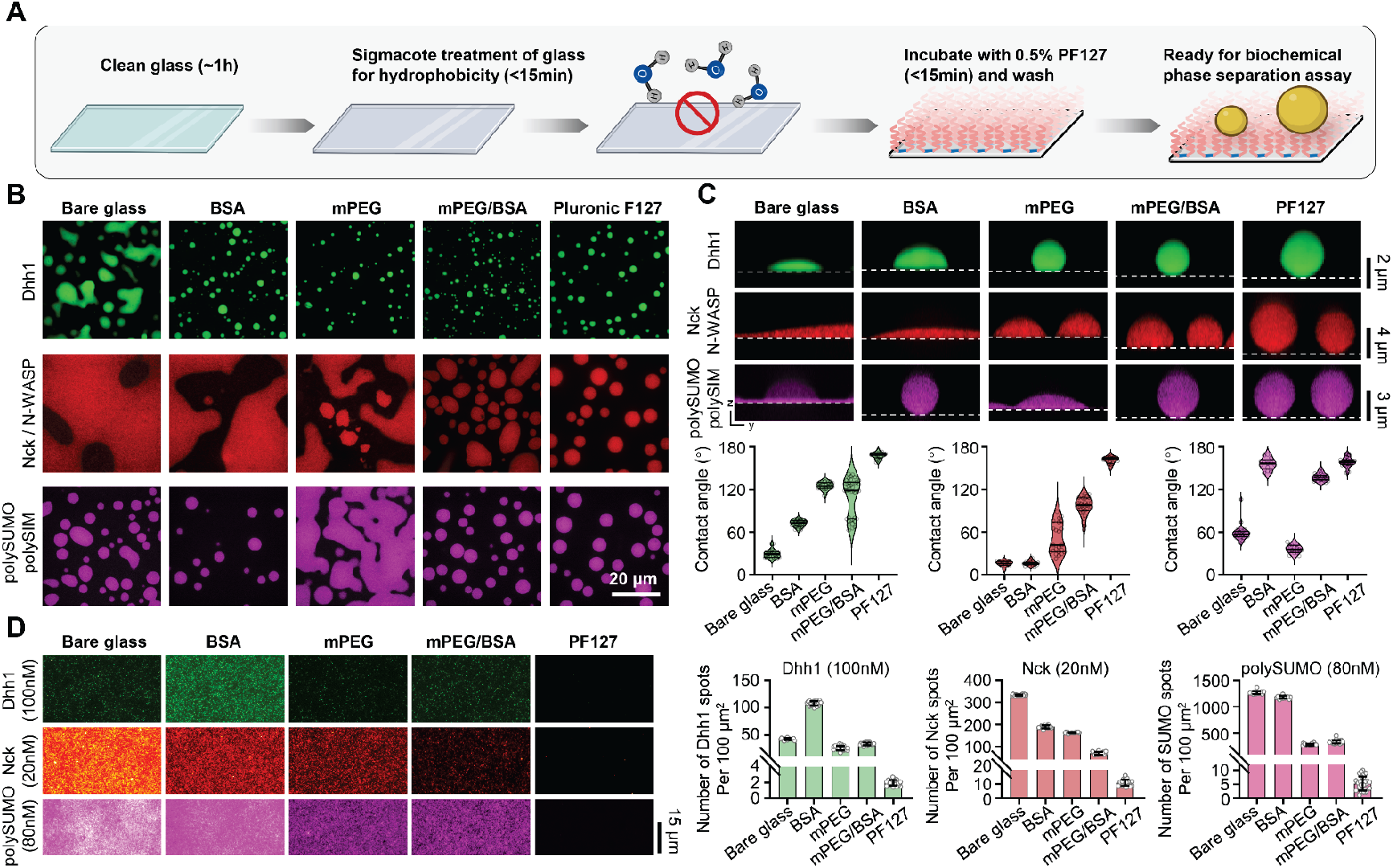
Self-assembly of PF127 as an efficient and broadly applicable surface passivation method for biochemical studies of biomolecular condensates. A, Workflow illustrating PF127 passivation. Image created with BioRender.com. B, Representative images showing Dhh1, Nck/N-WASP, and polySUMO/polySIM condensates on different passivated surfaces. C, Comparison of PF127 passivation against traditional techniques. Plots show the contact angle (median and quartiles, n>15 condensates per technique). D, Comparison of PF127 passivation with traditional techniques at single-molecule level (mean ± SD, n>20 images for each technique). Images for the same protein were captured under identical microscope settings and displayed with consistent brightness and contrast settings.

Relatively high concentrations (10-1000 nM) of free molecules often remain in the dilute phase surrounding condensates. These molecules can nonspecifically bind to slide surfaces, increasing background that impacts sensitive studies such as single-molecule imaging. To further evaluate the performance of PF127 passivation, we conducted single-molecule imaging of fluorescently labeled Dhh1, Nck and polySUMO at concentrations mirroring the dilute phase of these systems (tens of nM) (6, 7, 17). PF127 passivation continued to demonstrate superior performance in minimizing nonspecific binding compared to traditional methods (Fig. 2D). Thus, for both condensates and single-molecules, PF127 passivation appears to provide strong and generic blocking of surfaces.

### PF127 passivation is robust across diverse conditions

Next, we evaluated the robustness of PF127 treatment under diverse conditions: varying wash volumes, different pH values, and a spectrum of salt concentrations (Fig. 3A). We leveraged the strong wettability of Nck/N-WASP condensates (Fig. 2B and 2C) to monitor the impact of different treatments on the integrity of passivation. Based on condensate-surface contact angles, PF127 exhibits remarkable stability. It effectively resisted 10 ml of buffer wash on a 3.3×3.3 mm surface area (Fig. 3B) and remained stable in washes of pH 4-9 (Fig. 3C) or 0-1 M salt prior to imaging the condensates under normal buffer conditions (Fig. 3D). Furthermore, imaging millimeter-scale fields of view revealed excellent homogeneity of passivation across large areas (SI Appendix, Fig. S2). Finally, PF127 treated surfaces displayed no auto-fluorescence in commonly used excitation wavelengths of 430-790 nm (SI Appendix, Fig. S3).

**Figure 3.**
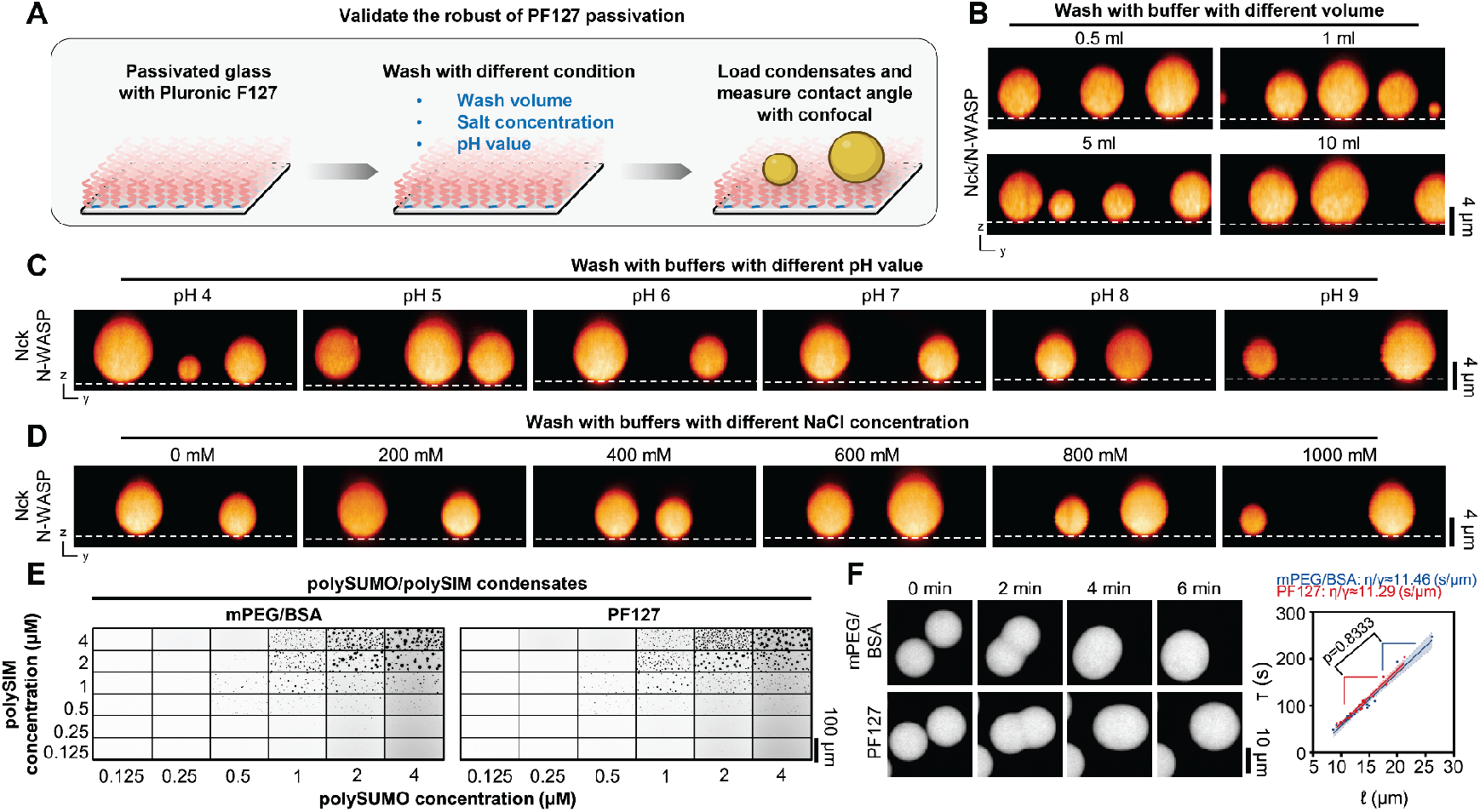
Validation of PF127 passivation robustness. A, Workflow for robustness validation of PF127 passivation. Image created with BioRender.com. B, Orthogonal views of Nck/N-WASP condensates on PF127-passivated surface after different buffer volume washes. Note: Imaging chamber area is 3.3x3.3 mm. C, Orthogonal views of Nck/N-WASP condensates on PF127-passivated surface pre-washed with buffers of pH values ranging from 4 to 9. The condensates were imaged under normal buffer conditions. D, Orthogonal views of Nck/N-WASP condensates on PF127-passivated surface pre-washed with buffer of varying NaCl concentrations (0 mM to 1000 mM). The condensates were imaged under normal buffer conditions. E, Comparison of phase separation threshold of polySUMO/polySIM condensates on glass passivated with PF127 versus mPEG/BSA. F, Comparison of inverse capillary velocity of polySUMO/polySIM condensates on glass passivated with PF127 versus mPEG/BSA (mean ± 95% CI, two-tailed t-test comparing slopes, n=22 for PF127 and n=17 for mPEG/BSA).

Even though excess free PF127 is removed during experiment procedures (Fig. 2, Detailed Protocol in SI Appendix) and assembled PF127 is highly stable (Fig. 3A-3D), a nonnegligible concern is the potential alteration of the properties of biomolecular condensates by surfactant. To address this concern, we selected polySUMO/polySIM condensates, a system that exhibited high contact angles in both mPEG/BSA and Pluronic F127 passivation (Fig. 2B and 2C), to investigate the impact of PF127 passivation on various condensate properties. PolySUMO/polySIM condensates generated on glass passivated with PF127 or mPEG/BSA showed essentially identical phase separation thresholds (Fig. 3E) and material properties (inverse capillary velocity) (Fig. 3F). Chromatin condensates, which exhibit variable properties based on surface passivation (9), also demonstrated similar phase separation thresholds (SI Appendix, Fig. S4A) and molecular exchange rates (SI Appendix, Fig. S4B and S4C) when generated on PF127-passivated surfaces compared to those previously reported for mPEG/BSA surfaces (9, 20). Thus, PF127 surfaces do not aberrantly affect condensate physical properties.

### Biotin-NeutrAvidin anchor points enable biomolecular condensates to be immobilized

A variety of investigations, including 3D imaging, microrheology, single-molecule imaging and fluorescence recovery after photobleaching (FRAP), benefit from condensate immobilization. However, due to the supreme passivation ability of PF127, condensates tended to move and rotate freely across the slide surface (Movie S1). To restrain movement, we treated hydrophobic glass with 1μg/ml biotin-labeled BSA prior to PF127 assembly (Fig. 4A), to create a low density (∼11 molecules/100 µm^2^) of stable, specific anchor points on the surface (Fig. 4B). In the presence of 15 nM NeutrAvidin, polySUMO/polySIM condensates containing 1 % biotin-labeled polySUMO were completely arrested on this surface but only minimally wet it, showing contact angles of 161° ± 7° (Fig. 4C-4E, Movie S2; note that we have not yet determined the exact number of anchor points needed to effectively immobilize individual condensates). Condensate immobilization also showed no obvious effect on molecular exchange rates (Fig. 4E). Thus, PF127-treatment can produce high levels of surface passivation while simultaneously allowing condensate immobilization.

**Figure 4.**
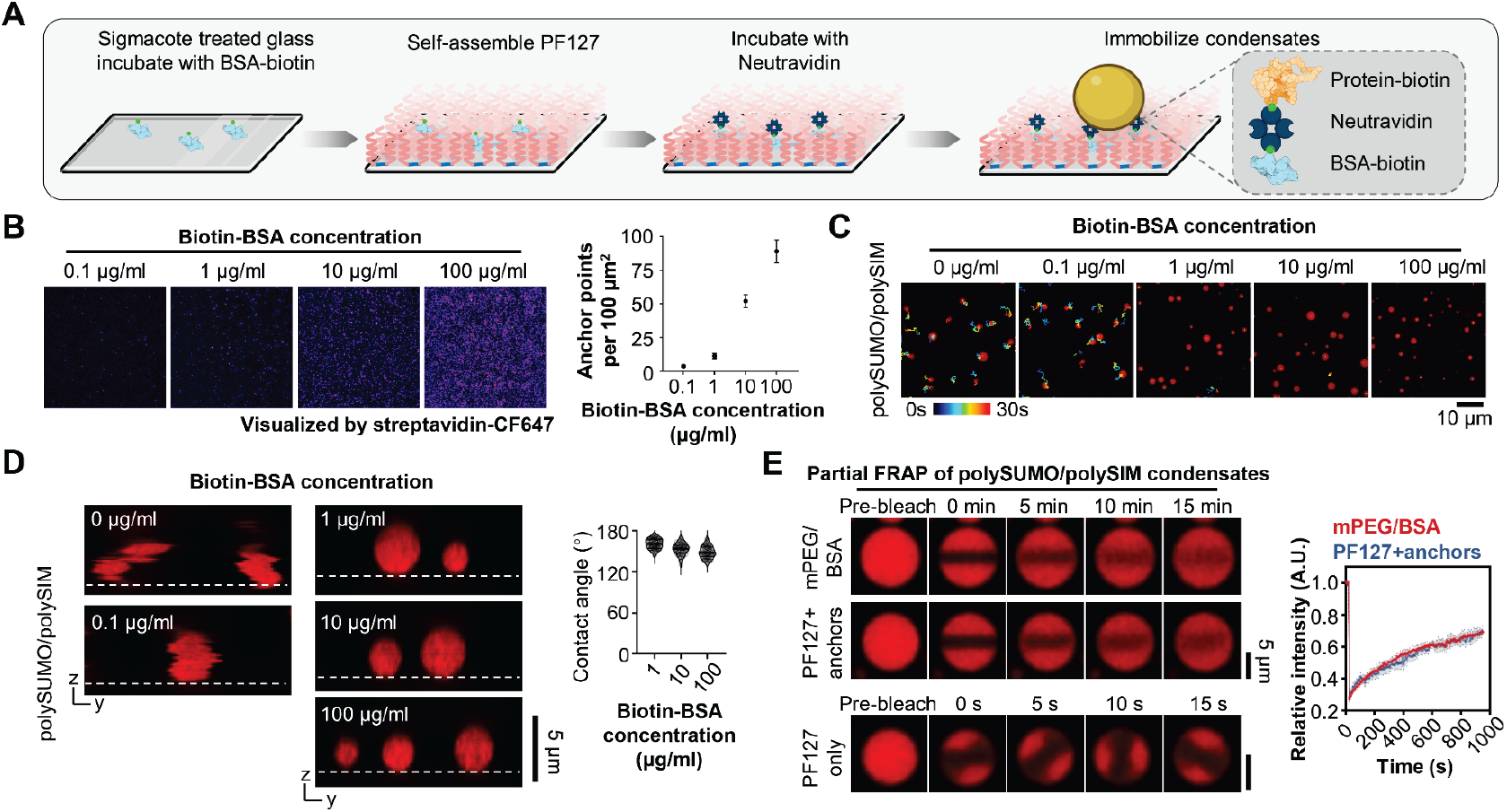
Immobilization of condensates through defined anchors. A, Workflow of condensates immobilization. Image created with BioRender.com. B, Representative TIRF imaging of anchor points on PF127-passivated glass incubated with varying concentrations of Biotin-BSA. Plot illustrates the density of anchor points for each concentration (mean ± SD, n>20 images for each concentration). C, Movement trajectories of polySUMO/polySIM condensates at varying biotin-BSA concentration. D, Orthogonal views of polySUMO/polySIM condensates at varying biotin-BSA concentration (note that apparently irregular shapes at low biotin-BSA concentrations are due to movement during imaging of the confocal stack). Plot illustrates contact angle of polySUMO/polySIM condensates at different biotin-BSA concentrations (median and quartiles, n>15 condensates for each condition). E, Partial FRAP of polySUMO/polySIM condensates on glass passivated by PF127 with anchor points or mPEG/BSA (mean ± SD, n=12 condensates for each condition). For comparison, partial FRAP of polySUMO/polySIM condensates on PF127-passivated glass without anchor points is displayed at a different time resolution at the bottom.

### Available polySIM binding sites are heterogeneously distributed within polySUMO/polySIM condensates

We then employed single-molecule imaging to visualize fluorescence recovery after photobleaching, monitoring exchange of individual fluorescent polySIM molecules between immobilized polySUMO/polySIM condensates and the dilute phase (Fig. 5A). Contrary to the slow overall exchange rate observed in traditional FRAP assays (Fig. 4E), single-molecule FRAP revealed the distinct appearance of individual fluorescence signals at the periphery of the condensates within hundreds of milliseconds after photobleaching (Fig. 5B). Comparisons between peripheral and central regions of polySUMO/polySIM condensates showed significantly faster recovery at the periphery (Fig. 5C). We first hypothesized that slow recovery at the center might be due to a low ability of new polySIM molecules to access the condensate core and slow diffusion from the periphery. However, single-molecule tracking showed that newly entered polySIM molecules readily and rapidly accessed all regions of the droplets (Fig. 5D, Movie S3). Analysis of the mobility patterns revealed that while polySIM molecules could access the center, many exhibited rapid diffusion there (Fig. 5D and 5E, Movie S4), with an appreciably larger mobile fraction compared to the periphery (Fig. 5F, Movie S5 and S6). These behaviors suggest that there may be a lower density of available binding sites capable of immobilizing polySIM molecules to the polymeric polySUMO/polySIM scaffold at the condensate center than at the periphery (Fig. 5G). Thus, availability of polySIM-binding sites, rather than accessibility, may account for the differing rates of fluorescence recovery and polySIM dynamics between the center and periphery. The higher polySIM recovery rate at peripheral regions may reflect surface tension, which could alter conformation, interactions, and potentially functions, of molecules within sub-condensate structures (21).

**Figure 5.**
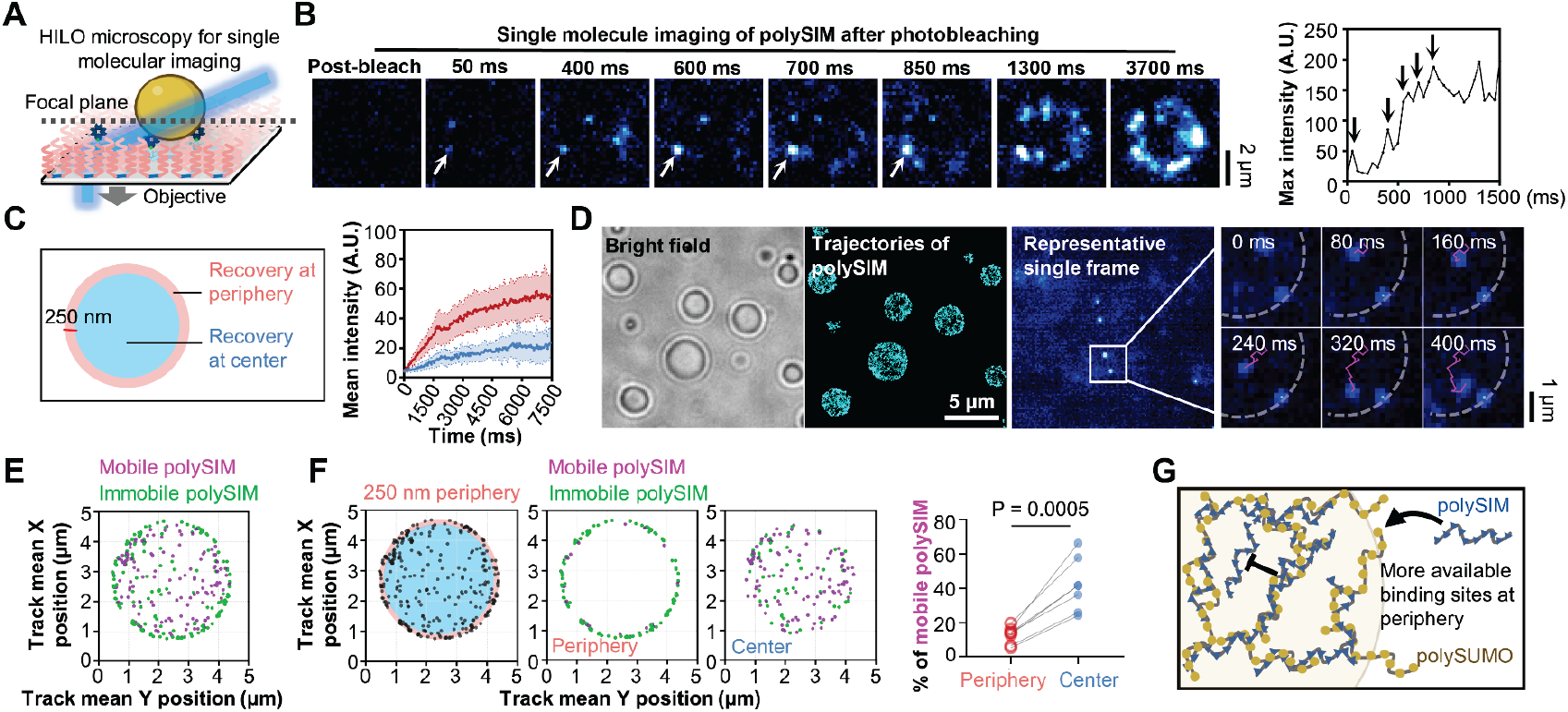
PF127 passivation facilitates high-precision single-molecule analyses. A, Schematic illustration of single-molecule imaging of immobilized condensates using highly inclined and laminated optical sheet (HILO) microscopy. Image created with BioRender.com. B, Single-molecule imaging of polySIM in polySUMO/polySIM condensate after photobleaching. Plot displays maximum intensity at the arrow-indicated spot. Vertical arrows within the plot mark time points at 50, 400, 600, 700, and 850 ms, correlating with the respective images on the left. C, Comparison of fluorescence recovery between the peripheral and central regions of condensates (mean ± SD, n=15 condensates for each condition). D, Trajectories of polySIM molecules within polySUMO/polySIM condensates (n>2000 tracks). Right image shows representative trajectories of polySIM molecules exhibiting varied mobility. Dashed line marks condensate position. The displayed area corresponds to the rectangle. E, Representative distribution pattern of mobile (average velocity > 3 μm/s, magenta) and immobile (average velocity > 3 μm/s, green) polySIM molecules within condensates. Mean X and Y positions of all spots in each track are shown. F, Proportions of immobile and mobile polySIM molecules at peripheral (250 nm from condensate surface) and central regions of the condensates (n=7 condensates, paired t-test). G, Potential explanation for fewer mobile molecules in peripheral regions.

## Discussion

An effective surface passivation method is crucial for preserving the material properties of biomolecular condensates, enabling accurate determination of their properties, and ensuring the success of biochemical studies. Traditional passivation methods, such as mPEG and BSA passivation, often do not sufficiently prevent nonspecific binding at concentrations used for *in vitro* reconstitution assays (10). This limitation makes those traditional methods suboptimal for various condensate systems, particularly those with high wettability, and fails to block adhesion of molecules to surfaces surrounding condensates, which can generate appreciable background in sensitive studies such as single-molecule imaging.

Our study introduces an efficient, robust, straightforward, and generally applicable surface passivation method via self-assembled PF127 for biochemical studies of biomolecular condensates. The method provides generic passivation for both condensates and surrounding molecules in dilute phases across diverse condensate systems, including IDR-mediated and multidomain-mediated phase separation systems with varying wettability, as well as highly sensitive chromatin condensates (Fig. 1 and 2). The method also demonstrates high resistance to extensive washing, wide ranges of pH 4-9, salt concentration 0-1 M, and ensures exceptional homogeneity over large areas with no auto-fluorescence background while preserving condensate properties (Fig. 3). When combined with a Biotin-NeutrAvidin system, the method allows condensate immobilization through specific anchor points while simultaneously producing high levels of surface passivation (Fig. 4). This facilitates multiple movement-sensitive methods that examine the bulk properties of biomolecular condensates, including 3D imaging, microrheology, FRAP, fluorescence correlation spectroscopy (FCS) and droplet fusion assays (22). This advance also paves the way for exploring molecular interactions within reconstituted condensates at the single-molecule level (Fig. 5).

Current surface passivation methods for biomolecular condensates involve complex, time-consuming experimental procedures, sometimes requiring hazardous chemical treatments (11, 12, 23) or specialized chemical knowledge (13, 24), which diminishes their accessibility and reproducibility. In contrast, PF127 passivation is straightforward, cost-effective, and requires less than one hour of active handling. The entire procedures can be completed within three hours without any additional experimental expertise, offering a 99% cost reduction compared to mPEG/BSA passivation and making it feasible for any standard biochemistry laboratory. The simplicity and efficiency of the method could enhance the throughput of experiments and increase accessibility of biochemical phase separation studies to a broader scientific community.

Our method employs PF127 self-assembly to prevent nonspecific biomolecule binding and condensate wetting, necessitating only the formation of the PF127 layer on hydrophobic surfaces to create a dense, brush-like structure without covalent interactions (15, 18, 25). Thus, it can potentially passivate other experimental materials, provided they are hydrophobic, expanding its application beyond microscopic observation and reducing experimental errors caused by nonspecific binding. Furthermore, as many surfactants can self-assemble on hydrophobic surfaces (26, 27), different surfactants could create surfaces with unique properties, which could facilitate studying material properties such as surface tension, hydrophobicity, and viscoelasticity of biomolecular condensates. In future developments, mixing different surfactants might generate a rough, heterogeneous surface, which could further create a super-repellent surface for biomolecular condensates studies (28).

## Materials and Methods

### Protein expression, purification, and conjugation

### Nck purification

BL21(DE3) cells expressing GST-Nck were collected by centrifugation and lysed by sonication in 25 mM Tris-HCl (pH 8.0), 200 mM NaCl, 2 mM EDTA (pH 8.0), 1 mM DTT, 1 mM PMSF, 1 μg/ml antipain, 1 μg/ml benzamidine, 1 μg/ml leupeptin, and 1 μg/ml pepstatin. Proteins were affinity-purified with Glutathione Sepharose 4B (GE Healthcare). GST was cleaved from protein by TEV protease treatment for 16 hours at 4 °C, followed by anion exchange chromatography using Source 15 Q resin. Eluted protein was pooled and purified further by a Source 15 S cation exchange column. Eluted protein was concentrated using Amicon Ultra Centrifugal Filter units (Millipore) and further purified by size exclusion chromatography using a Superdex 75 prepgrade column (GE Healthcare) in 25 mM HEPES (pH 7.5), 150 mM NaCl, and 1 mM DTT.

### N-WASP purification

BL21(DE3) cells expressing His6-N-WASP were collected by centrifugation and lysed by cell disruption (Emulsiflex-C5, Avestin) in 20 mM imidazole (pH 7.0), 300 mM KCl, 5 mM βME, 0.01% NP-40, 1 mM PMSF, 100 μM antipain, 1 mM benzamidine, 100 μM leupeptin, and 1 μM pepstatin. Proteins were affinity-purified with NiNTA agarose (Qiagen). The eluate was further purified over a Source 15 Q column (Cytiva). The His6-tag was removed by TEV protease treatment at 4 °C for 16 hours. Cleaved N-WASP was then applied to a Source 15 S column (Cytiva). Fractions containing N-WASP were concentrated using Amicon Ultra Centrifugal Filter units (Millipore) and further purified by size exclusion chromatography using a Superdex 200 prepgrade column (Cytiva) in 25 mM HEPES (pH 7.5), 150 mM KCl, 1 mM DTT, and 10% glycerol.

### PolySUMO purification

BL21(DE3) cells expressing His6-PolySUMO were collected by centrifugation and lysed by cell disruption (Emulsiflex-C5, Avestin) in 20 mM imidazole (pH 7.0), 300 mM KCl, 5 mM βME, 1 mM PMSF, 100 μM antipain, 1 mM benzamidine, 100 μM leupeptin, and 1 μM pepstatin. Proteins were affinity-purified with Ni-NTA Agarose Resin (Qiagen), followed by cation exchange chromatography using Source 15S Resin (Cytiva). Purified fractions were pooled and cleaved with TEV protease and flowed through Source 15S resin to remove un-cleaved proteins. Cleaved protein products were purified further by anion exchange chromatography using Source 15Q Resin (Cytiva), followed by size exclusion chromatography using a Superdex 200 gel filtration column (Cytiva) in 25 mM HEPES (pH 7.5), 150 mM KCl, 1 mM DTT, and 10% glycerol.

### PolySIM purification

BL21(DE3) cells expressing His6-PolySIM were collected by centrifugation and lysed by cell disruption (Emulsiflex-C5, Avestin) in 20 mM imidazole (pH 7.0), 300 mM KCl, 5 mM βME, 1 mM PMSF, 100 μM antipain, 1 mM benzamidine, 100 μM leupeptin, and 1 μM pepstatin. Proteins were affinity-purified with Ni-NTA Agarose Resin (Qiagen), followed by cation exchange chromatography using Source 15S Resin (Cytiva). Purified fractions were pooled and cleaved with TEV protease and flowed through Source 15S resin to remove un-cleaved proteins. Cleaved protein products were purified further by anion exchange chromatography using Source 15Q Resin (Cytiva), followed by size exclusion chromatography using a Superdex 200 gel filtration column (Cytiva) in 25 mM HEPES (pH 7.5), 150 mM KCl, 1 mM DTT, and 10% glycerol.

### His6-MBP-Dhh1-mEGFP purification

BL21(DE3) cells expressing His6-MBP-mEGFP-Dhh1 were collected by centrifugation and lysed by cell disruption (Emulsiflex-C5, Avestin) in 20 mM Tris (pH 8.0), 500 mM NaCl, 10% glycerol, 10 mM imidazole, 5 mM βME. Proteins were affinity-purified with Ni-NTA Agarose Resin (Qiagen), followed by affinity-purification with amylose resin (New England Biolabs). The amylose eluates were purified further by anion exchange chromatography using Source 15Q Resin (Cytiva), followed by size exclusion chromatography using a Superdex 200 gel filtration column (Cytiva) in 10 mM MES (pH 7), 300 mM KOAc, 5 mM BME, and 5% glycerol.

### Conjugation

Recombinant proteins to be labeled with fluorophores were concentrated using Amicon Ultra Centrifugal Filter units (Millipore) to ∼100 μM. 5 mM βME was added to reduce cysteine residues followed by buffer exchange using a HiTrap 26/10 Desalting column (GE Healthcare) in 25 mM HEPES (pH 7.5) and 150 mM NaCl. Fractions containing protein were collected and concentrated to 100 μM. 500 μM Alexa Fluor 647 C2 Maleimide (ThermoFisher, for polySUMO), Atto 647N Maleimide (Sigma, for Nck and polySIM) or Biton Maleimide (Sigma, for polySUMO) was added, and the reaction was incubated with gentle mixing at 4 °C for 16 hours. The reaction was quenched with 1 μl 14.3 M βME followed by final buffer exchange using size exclusion chromatography (GE Healthcare).

Polynucleosomal arrays were purified, assembled and conjugated as previously described (20).

### Preparation of PF127 passivated surface

#1.5H glass slides (Thorlabs) or #1.5H 384-well microscopy plates (Cellvis) were cleaned using a 1-hour wash with 5% Hellmanex and thorough rinsing with H2O, followed by a 1-hour etching with 1M KOH and another copious rinse with H2O (See Supplementary Protocol for detailed procedures). Subsequently, the glass surfaces were treated with Sigmacote (Sigma) according to the manufacturer’s instructions. The slides or plates were then rinsed with isopropanol and allowed to dry completely in a chemical hood and can be stored at this stage for up to several months.

For passivation without anchor points, slides or plates were then incubated with 0.5% PF127 (Sigma) in 20 mM Tris (pH8.0), 150 mM KCl for 15 min at room temperature. A final wash with the phase separation assay buffer was performed twice. The slides or plates were then ready for use. It is important that once treated with PF127 the slides or plates must be kept under an aqueous solution; drying irreversibly damages the self-assembled surfactant layer.

For passivation with anchor points, slides or plates were incubated with 1-10 μg/ml biotinylated BSA (Sigma) in 20 mM Tris (pH 8.0), 500 mM KCl for 5 min. This step was followed by a 15-minute incubation with 0.5% PF127 (Sigma) in 20 mM Tris (pH 8.0), 150 mM KCl at room temperature. Afterward, the slides or plates were washed twice with 20 mM Tris (pH 8.0), 150 mM KCl and incubated with 15 nM NeutrAvidin in the same buffer for 5 minutes. A final wash with the phase separation assay buffer was performed twice. The slides or plates were then ready for use. A detailed protocol is available (SI Appendix).

### Preparation of mPEG/BSA passivated surface

mPEG passivation was performed as previously described (9). Briefly, #1.5H glass slides (Thorlabs) or #1.5H 384-well microscopy plates (Cellvis) were washed with 5% Hellmanex at 37°C for 4 hours and then rinsed copiously with H2O. Silica was etched with 1 M NaOH for 1 hour at room temperature and then rinsed copiously with H2O. Depolymerized Silica was covalently bonded overnight at room temperature to 20 mg/mL 5K mPEG-silane (PEGWorks) suspended in 95% Ethanol. The slides or plates were washed many times with 95% Ethanol, rinsed with copious amounts of H2O, and completely dried in a chemical hood over 3-4 hours. The slides or plates were rinsed three times with phase separation assay buffer and were then ready for use. For mPEG/BSA passivation, PEGylated glass was further incubation with freshly prepared 100 mg/mL BSA in 20 mM Tris (pH 8.0), 500 mM KCl for 30 minutes. The slides or plates were rinsed three times with phase separation assay buffer to remove BSA and were then ready for use.

For BSA passivation, clean and etched glass was directed incubation with freshly prepared 100 mg/mL BSA in 20 mM Tris (pH 8.0), 500 mM KCl for 30 minutes. The slides or plates were rinsed three times with phase separation assay buffer to remove BSA and were then ready for use.

### Biomolecular condensates reconstitution

Nck/N-WASP condensates reconstitution: Nck (containing 10% Atto 647N-labeled Nck) and N-WASP proteins were mixed in 25 mM HEPES (pH 7.5), 150 mM KCl to achieve a final concentration of 10 μM, unless otherwise noted, and incubated for 1 hour at room temperature before imaging. PolySUMO/polySIM condensate reconstitution: polySUMO (containing 1% Biotin labeled polySUMO in immobilization assays) and polySIM (containing 10% Atto 647N-labeled polySIM for conventional imaging, 1% Atto 647N-labeled polySIM for single-molecule imaging after photobleaching and 0.1% Atto 647N-labeled polySIM for single-molecule tracking) proteins were mixed in 25 mM HEPES (pH 7.5), 150 mM KCl to achieve a final concentration of 5 μM concentration unless otherwise note, and incubated for 1 hour at room temperature before imaging. Dhh1 condensate reconstitution: His6-MBP-mEGFP-Dhh1 was diluted in 10 mM MES (pH 6), 50 mM KOAc buffer, and TEV protease was added at a 1:50 molar ratio to remove the His6-MBP tag and initiate condensate formation. The reaction mixture was incubated for 6 hours at room temperature before imaging.

Chromatin condensate reconstitution: Polynucleosomal arrays, featuring either 25 bp or 30 bp inter-nucleosome linker DNA and 1% Alexa 594-labeled histone H2B, were diluted in 25 mM Tris-Acetate (pH 7.5), 150 mM KOAc and 1 mM Mg(OAc)2 buffer to a final concentration of 1 μM. The mixture was incubated for 1 hour at room temperature before imaging.

### Microscopy

Spinning disk images were captured using a Leica DMI6000 microscope, equipped with a Yokogawa CSU-X1 spinning disk confocal scanner unit and a Hamamatsu ImagEMX2 EM-CCD camera. For high resolution images, a Leica 100× 1.49 NA oil immersion objective was utilized (as shown in Fig. 1B, 1C, 2B, 2C. 3B-3D, 4C, 4D, Appendix Fig. S4A). Broader field views were obtained using a Leica 20× 0.4 NA air objective (as shown in Fig. 3E and 3F).

Confocal images were acquired on a Leica SP8 microscope equipped with a 63 × 1.4 NA oil immersion objective. For detection, the system employed either hybrid (HyD) detectors (SI Appendix Fig. S2 and S3) or PMT detectors (Fig. 4E).

Total Internal Reflection Fluorescence (TIRF) images were captured using a TIRF/iLAS2 TIRF/FRAP module (Biovision) mounted on a Leica DMI6000 microscope base equipped with Leica 100× 1.49 NA oil immersion objective (Fig. 4B) or a DeltaVision OMX SR system, equipped with an Olympus 60 × 1.49 NA oil immersion objective for TIRF and ring-TIRF capabilities (Fig. 2D).

Highly inclined and laminated optical sheet (HILO) microscopy for single molecule imaging was captured using DeltaVision OMX SR system, equipped with an Olympus 60 × 1.49 NA oil immersion objective for TIRF (Fig. 5B-5F).

### Measuring contact angle

To measure the contact angles, stacks of condensates were captured using a spinning disk confocal microscope with 100× objective. Spherical aberration artifacts were corrected as described previously (29). These image stacks were then processed using Fiji/ImageJ software to generate orthogonal (y-z) views of the condensates. The contact angles were subsequently measured utilizing the angle tool in Fiji/ImageJ.

### Measuring inverse capillary velocity

Time-lapse imaging of condensate fusion events was captured using a spinning disk confocal microscope equipped with 20× objective, capturing frames at intervals of 4.47 s. We selected fusion events involving two droplets of comparable size for analysis. The method for determining inverse capillary velocity was adapted from previously published protocols (30).

In brief, the aspect ratio (A.R.) of the condensates was calculated by fitting an ellipse to the shape and computing A.R. = *𝓁*long/*𝓁*short, where *𝓁*long and *𝓁*short represent the long and short axes of the ellipse, respectively. The time evolution of this aspect ratio was fit using the function A.R. = 1 + (A.R.0 - 1) × exp(-t/τ), where t is time, τ is the characteristic relaxation time, and A.R.0 is the initial aspect ratio.

For these fusing condensates, we defined the length scale as the geometric mean *𝓁* = [(*𝓁*long(t = 0) - *𝓁*short(t = 0)) × *𝓁*short(t = 0)]1/2. The relationship between τ and *𝓁* was characterized by fitting the data to a linear equation τ = (η/γ) × *𝓁*, from which we determined the inverse capillary velocity, η/γ.

### Validation of PF127 passivation robustness

To validate the robustness of PF127 passivation, #1.5H 384-well microscopy plates (Cellvis) were first passivated by PF127 as described above. The wells then underwent various treatments to test the stability of the passivation:

1.Washing with varying volumes (0.5, 1, 5, and 10 ml) of 20 mM Tris (pH 8.0), 150 mM KCl buffer.

2.Incubation with 100 μl of different buffers for 10 minutes at room temperature: 20 mM MES (pH 4.0), 20 mM MES (pH 5.0), 20 mM MES (pH 6.0), 20 mM HEPES (pH 7.0), 20 mM Tris (pH 8.0), and 20 mM Tris (pH 9.0).

3.Incubation with 100 μl of 20 mM HEPES (pH 7.5) containing varying concentrations of NaCl (0, 200, 400, 600, 800, and 1000 mM) for 10 minutes at room temperature.

Following these treatments, the wells were washed twice with the buffer used for condensate reconstitution. Condensates were then loaded into the wells for imaging.

### Characterization of auto-fluorescent of PF127 passivation

To evaluate the auto-fluorescent properties of PF127 passivation, #1.5H 384-well microscopy plates (Cellvis) were first passivated by PF127 as described above. After passivation, each well was filled with 50 μl of 0.5% PF127 solution for imaging purposes. For a bare glass control, 50 μl of water was added directly into untreated wells. Imaging was conducted using a Leica SP8 microscope using the λ scanning function. Excitation was performed using 405 nm, 488 nm, 552 nm and 638 nm lasers, while emission light ranging from 430 to 790 nm was collected in 20 nm bandwidth increments.

### Fluorescence recovery after photobleaching (FRAP)

FRAP assays were perform using a Leica SP8 microscope equipped with a 100 × 1.4 NA oil immersion objective (Fig. 2c) or a DeltaVision OMX SR system equipped with a 60 × 1.42 NA oil immersion objective and ring-TIRF capabilities (Extended Data Fig. 6b).

In these assays, a designated region of interest (ROI) was subjected to photobleaching, and the subsequent recovery of fluorescence intensity within the ROI was monitored and recorded for each experiment. The intensity recovery curves were then normalized and corrected for photobleaching effects, following methodologies described previously (31). Briefly, the fluorescence signal measured in a region of interest normalized to the change in total fluorescence was determined as I = (It/I0) × (T0/Tt), where I0 is the average intensity of the unbleached region of interest at time point t. I0 is the average pre-bleach intensity of the region of interest. T0 and Tt are the average fluorescence intensities of a neighboring condensate in the same field of view at pre-bleach or at timepoint t, respectively. The recovery curves were fit to the following expression using GraphPad as Y(t) = A×(1-e^(τ×t)), where A is the end-value of the recovered intensity, τ is the fitted parameter and t is the time after the bleaching.

### Single molecular tracking and analysis

Single molecule imaging was performed using a DeltaVision OMX SR system, equipped with a 60 × 1.49 NA TIRF oil immersion objective and ring-TIRF system. Imaging was conducted in epi/TIRF mode using CMOS cameras set to 2×2 binning and a beam condenser to enhance contrast. For single molecule imaging after photobleaching (refer to Fig. 2e and 2f), the photokinetic mode was employed to bleach the condensates, and signal recovery was captured at a time resolution of 50 ms. A total of approximately 150 images (∼7.5 seconds) were recorded for each event. The images were processed using Fiji/ImageJ for background subtraction. The mean intensity in different regions of the condensates (periphery and center) was measured and plotted.

For single molecule tracking of newly entered polySIM (refer to Fig. 2h and 2i), the burst mode was used at 50 Hz imaging rate. Movies were first processed using Fiji/ImageJ for background subtraction. The first 100 frames of each movie were discarded to ensure that only newly entered polySIM molecules were imaged. The movies were then analyzed using TrackMate in Fiji/ImageJ. Specifically, spots were detected using the Hessian detector with a 550 nm diameter and sub-pixel localization. Spots were then filtered based on contrast, quality, signal-to-noise ratio, and maximum intensity. Track generation was accomplished using the LAP Tracker with a maximum frame-to-frame linking distance of 500 nm and a maximum gap of 2 frames, also at a maximum distance of 500 nm. Tracks consisting of at least 6 events were retained, and trajectories were plotted. The average velocity of tracks was used to categorize the events: tracks with average velocity < 3 μm/s were classified as immobile, while those > 3 μm/s were classified as mobile. The mean of the X and Y position of all the spots in each track was used to indicate the position of polySIM molecules. The number of polySIM in each mobility category was counted in both the peripheral and central regions of the condensates.

## Acknowledgments

We thank June Ho Hwang for providing the mEGFP-Dhh1 protein, Sabareesan A. Thody for providing the polySUMO and polySIM protein, Huabin Zhou for providing the polynucleosomal arrays, Xuemeng Zhen for testing the PF127 passivation, and Rosen lab members for discussion. We would like to acknowledge the Quantitative Light Microscopy Core, a Shared Resource of the Harold C. Simmons Cancer Center, supported in part by an NCI Cancer Center Support Grant, 1P30 CA142543-0. Research was supported by the Howard Hughes Medical Institute, the Mar Nell and F. Andrew Bell Chair in Biochemistry (to M.K.R.) and the Welch Foundation (I-1544 to M.K.R.). This article is subject to HHMI’s Open Access to Publications policy. HHMI lab heads have previously granted a nonexclusive CC BY 4.0 license to the public and a sublicensable license to HHMI in their research articles. Pursuant to those licenses, the author-accepted manuscript of this article can be made freely available under a CC BY 4.0 license immediately upon publication.

## References

1. S. F. Banani, H. O. Lee, A. A. Hyman, M. K. Rosen, Biomolecular condensates: organizers of cellular biochemistry. Nat Rev Mol Cell Biol 18, 285–298 (2017).

2. Y. Shin, C. P. Brangwynne, Liquid phase condensation in cell physiology and disease. Science 357 (2017).

3. A. S. Lyon, W. B. Peeples, M. K. Rosen, A framework for understanding the functions of biomolecular condensates across scales. Nat Rev Mol Cell Biol 22, 215–235 (2021).

4. S. Alberti et al., A User’s Guide for Phase Separation Assays with Purified Proteins. J Mol Biol 430, 4806–4820 (2018).

5. D. M. Mitrea et al., Methods for Physical Characterization of Phase-Separated Bodies and Membrane-less Organelles. J Mol Biol 430, 4773–4805 (2018).

6. P. Li et al., Phase transitions in the assembly of multivalent signalling proteins. Nature 483, 336–340 (2012).

7. S. L. Currie et al., Quantitative reconstitution of yeast RNA processing bodies. Proc Natl Acad Sci U S A 120, e2214064120 (2023).

8. S. Alberti, A. Gladfelter, T. Mittag, Considerations and Challenges in Studying Liquid-Liquid Phase Separation and Biomolecular Condensates. Cell 176, 419–434 (2019).

9. B. A. Gibson et al., In diverse conditions, intrinsic chromatin condensates have liquid-like material properties. Proc Natl Acad Sci U S A 120, e2218085120 (2023).

10. M. L. Visnapuu, D. Duzdevich, E. C. Greene, The importance of surfaces in single-molecule bioscience. Mol Biosyst 4, 394–403 (2008).

11. S. D. Chandradoss et al., Surface passivation for single-molecule protein studies. J Vis Exp 10.3791/50549 (2014).

12. S. Shen, M. Naganuma, Y. Tomari, H. Tadakuma, Revisiting the Glass Treatment for Single-Molecule Analysis of ncRNA Function. Methods Mol Biol 2509, 209–231 (2022).

13. A. Testa et al., Surface Passivation Method for the Super-repellence of Aqueous Macromolecular Condensates. Langmuir 39, 14626–14637 (2023).

14. S. Ghosh, A. Ray, N. Pramanik, Self-assembly of surfactants: An overview on general aspects of amphiphiles. Biophys Chem 265, 106429 (2020).

15. M. W. H. Kirkness, C. S. Korosec, N. R. Forde, Modified Pluronic F127 Surface for Bioconjugation and Blocking Nonspecific Adsorption of Microspheres and Biomacromolecules. Langmuir 34, 13550–13557 (2018).

16. B. Hua et al., An improved surface passivation method for single-molecule studies. Nat Methods 11, 1233–1236 (2014).

17. S. F. Banani et al., Compositional Control of Phase-Separated Cellular Bodies. Cell 166, 651–663 (2016).

18. M. R. Nejadnik et al., Adsorption of pluronic F-127 on surfaces with different hydrophobicities probed by quartz crystal microbalance with dissipation. Langmuir 25, 6245–6249 (2009).

19. L. Shen, A. Guo, X. Zhu, Tween surfactants: Adsorption, self-organization, and protein resistance. Surface Science 605, 494–499 (2011).

20. B. A. Gibson et al., Organization of Chromatin by Intrinsic and Regulated Phase Separation. Cell 179, 470–484 e421 (2019).

21. M. Farag et al., Condensates formed by prion-like low-complexity domains have small-world network structures and interfaces defined by expanded conformations. Nat Commun 13, 7722 (2022).

22. L. R. Ganser, S. Myong, Methods to Study Phase-Separated Condensates and the Underlying Molecular Interactions. Trends Biochem Sci 45, 1004–1005 (2020).

23. C. He, C. Y. Wu, W. Li, K. Xu, Multidimensional Super-Resolution Microscopy Unveils Nanoscale Surface Aggregates in the Aging of FUS Condensates. J Am Chem Soc 145, 24240–24248 (2023).

24. A. Hucknall, S. Rangarajan, A. Chilkoti, In pursuit of zero: polymer brushes that resist the adsorption of proteins. Advanced Materials 21, 2441–2446 (2009).

25. C. Wu, G. Xia, J. Sun, R. Song, Synthesis and anisotropic self-assembly of Ag nanoparticles immobilized by the Pluronic F127 triblock copolymer for colorimetric detection of H 2 O 2. RSC advances 5, 97648–97657 (2015).

26. C. Schroen, M. C. Stuart, K. Van der Voort Maarschalk, A. Van der Padt, K. Van’t Riet, Influence of preadsorbed block copolymers on protein adsorption: surface properties, layer thickness, and surface coverage. Langmuir 11, 3068–3074 (1995).

27. G. B. Sigal, M. Mrksich, G. M. Whitesides, Using surface plasmon resonance spectroscopy to measure the association of detergents with self-assembled monolayers of hexadecanethiolate on gold. Langmuir 13, 2749–2755 (1997).

28. A. Cassie, S. Baxter, Wettability of porous surfaces. Transactions of the Faraday society 40, 546–551 (1944).

29. E. E. Diel, J. W. Lichtman, D. S. Richardson, Tutorial: avoiding and correcting sample-induced spherical aberration artifacts in 3D fluorescence microscopy. Nat Protoc 15, 2773–2784 (2020).

30. C. P. Brangwynne, T. J. Mitchison, A. A. Hyman, Active liquid-like behavior of nucleoli determines their size and shape in Xenopus laevis oocytes. Proc Natl Acad Sci U S A 108, 4334–4339 (2011).

31. R. D. Phair, S. A. Gorski, T. Misteli, Measurement of dynamic protein binding to chromatin in vivo, using photobleaching microscopy. Methods Enzymol 375, 393–414 (2004).

