## Supplementary material for "Advanced Surface Passivation for High-Sensitivity Studies of Biomolecular Condensates": SI Appendix

**This PDF file includes:**

Supporting text: detailed Protocol of PF127 passivation
Figures S1 to S4
Legends for Movies S1 to S6

**Other supporting materials for this manuscript include the following:**

Movies S1 to S6

### Supporting Information Text

#### Detailed protocol to passivate glass surface by Pluronic F127 self-assembly for in vitro biomolecular condensate studies

#### Reagents

For all subsequent preparations outlined in this protocol, 'MilliQ H2O' refers to water with a resistivity of at least 18.2 MΩ cm and a total organic carbon (TOC) content of no more than 5 ppm.

- Hellmanex™ III Liquid Cleaning Concentrate (Fisher Scientific, 14-385-864)
- 384 Well glass bottom plate with high performance #1.5 cover glass (Cellvis, P384-1.5H-N)
- MilliQ $H_2O$
- Potassium hydroxide (Sigma-Aldrich, 221473)
- Thermo Scientific™ Adhesive PCR Foil Seal (Thermo Scientific, AB0626)
- Sigmacote (Sigma-Aldrich, SL2)
- Pluronic F127 (Sigma-Aldrich, P2443)
- Isopropanol (Fisher Scientific A416-4)
- NeutrAvidin (Thermo Scientific, 31000)
- Biotin-BSA (Sigma-Aldrich, A8549)

#### Equipment

- Fume hood
- Microwave oven
- Glass syringe (We didn't test plastic syringe)

#### Procedure

1. Cleaning the 384-well glass-bottom plate

- Prepare 1 L of 5 % Hellmanex solution or 2 % Alconox solution (diluted with MilliQ H2O) in a 1 L glass beaker.
- Gently heat the solution by microwaving for 1 minute on high power, aiming for approximately 50℃.
- Carefully remove the beaker from the microwave. Submerge the 384-well glass-bottom plate in the solution, ensuring the well side is facing up. This orientation allows the solution to fill each well completely, avoiding air pockets. Leave to soak for 30 minutes.
- Optional Step: After the initial 30-minute soak, remove the plate and gently tap it to dislodge any air bubbles on the glass surface. Re-submerge the plate in the solution with the well side up for another 30-minute soak. This ensures thorough cleaning of the plate's bottom.
- Remove the plate from the solution, being careful not to empty the wells. Place the plate on a clean surface. Rinse and refill the beaker with 1 liter of tap water. Submerge the plate again, maintaining the well side up orientation. Gently rotate the plate by hand for about 10 seconds to ensure thorough rinsing.
- Repeat the rinsing step 10 times with tap water.
- Repeat the rinsing step another 10 times, this time using MilliQ H2O.
- Discard water from 384-well plate.

Note: Cleaning the glass may not be necessary if the plates are pre-cleaned but highly recommended.

2. Etching glass with hydroxide

- Tap inverted plate on a clean paper towel to remove excess water.
- Fill each well with 100 µL 1 M KOH. Let it sit for 1 hour at room temperature.

- Place the plate on a clean table. Rinse the beaker, fill with 1L tap water, and submerge the plate, well side up. Rotate for 10 s.
- Repeat the rinsing step 10 times with tap water.
- Repeat the rinsing step another 10 times, this time using MilliQ H2O.
- Discard water from 384-well plate.

Note: Etching with KOH is optional but recommended for high precision experiments.

3. Siliconization using Sigmacote for hydrophobicity

- Perform all siliconization steps in a fume hood.
- Draw about 1 mL of Sigmacote into a glass syringe.

Note: If using a plastic syringe, ensure it's compatible with Sigmacote.

- Dispense about 100µL Sigmacote into the first well, then immediately drain. Continue for each well, ensuring complete coverage. The reaction is almost instantaneous so you can process this step very fast as long as Sigmacote covers glass surface.
- Allow the plate to air dry in the hood for about 10 min.

Note: Oil-like residues may appear but will be washed away in subsequent steps.

- Rinse the plate twice with isopropanol.
- Fully dry the plate in the hood.
- Cover the entire dried plate with PCR foil. The plate can be kept for at least a few months.

The following 4 and 5 sections describe two methods for assembling Pluronic F127. The first method, passivation without anchor points, is appropriate for routine phase separation assays, such as visualizing fusion or constructing phase diagrams. The second method, involving passivation with anchor points, is designed for immobilizing droplets during prolonged imaging processes like 3D imaging, FRAP, and single molecule tracking.

4. Assembling Pluronic F127 without Anchor Point

- Remove foil from required wells using a scalpel.
- Fill wells with 100 µL 0.5%(w/v) Pluronic F127 in buffer that will be used in phase separation assay and incubate for > 15min.
- After incubation, sequentially wash each well with buffer used in phase separation assay for at least 5 times (sequentially wash means first remove 90 µL buffer, then add 90 µL fresh buffer, then remove 90 µL buffer and repeat for at least 5 times. <u>DO NOT dry the glass at any time</u> or the assembled F127 will be destroyed. One can also wash the well by suck up the buffer using a vacuum system while adding 500 µL wash buffer by pipette).
- Discard the excess solution but leave ~5 µL buffer in the well. <u>DO NOT dry the well</u>.
- Note: Although drying the wells should generally be avoided, it has been observed that some types of condensates can maintain their integrity on dried glass.
- Wells are now ready for the phase separation assay.

5. Assembling Pluronic F127 with anchor point

- Remove foil from required wells using a scalpel.
- Incubate wells with 20 µL1-100 µg/ml Biotin-BSA in 50mM Tris (pH8), 500mM NaCl buffer for 5 minutes at room temperature.

Note: 10 µg/ml Biotin-BSA is a good start point for non-sticky condensates.

- Wash wells twice with 200µl 50mM Tris (pH8), 500mM NaCl buffer.
- Fill wells with 100 µL 0.5%(w/v) Pluronic F127 in buffer that will be used in phase separation assay. Incubate for > 15min.
- Sequentially wash each well with the buffer at least 5 times. <u>Avoid drying the glass</u>. (See section 4 for details)
- Incubate wells with 20 µL 15 nM NeutrAvidin in buffer that will be used in phase separation assay for 5 minutes at room temperature.
  Wash wells with 200µl buffer. <u>DO NOT dry the glass at any time</u>.
- Discard the excess solution but leave ~5 µL buffer in the well. <u>DO NOT dry the well</u>.

- Wells are now ready for the phase separation assay.

Note: Add a small amount of biotin-labeled phase separation protein when preparing your sample. About 1% biotin-labeled protein is usually sufficient to immobilize condensates with low wettability. For sticky condensates, Biotin-NeutrAvidin-Biotin interaction may not be necessary. Incubating with BSA before F127 assembly will be enough to create sticky regions for immobilizing condensates.

**Time Taken**

- Clean the 384-well glass-bottom plate: ~1 h
- Etch glass with hydroxide: ~1 h
- Siliconize glass: 15-30 min
- Assemble Pluronic F127: 20 min

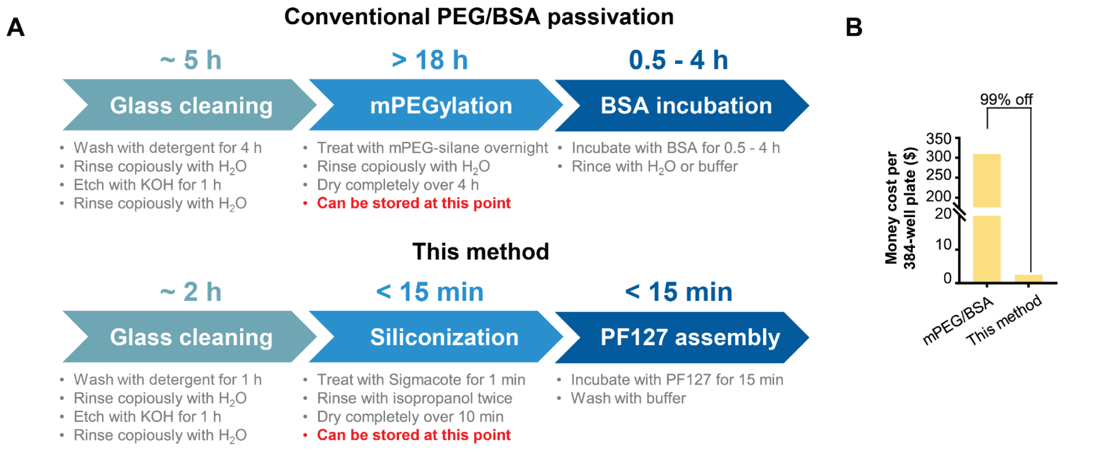


**Fig. S1. Comparative analysis of time and cost for PF127 and mPEG/BSA passivation.**

A, Workflow and time cost comparison between PF127 passivation and mPEG/BSA passivation. Image created with BioRender.com.

B, Comparison of financial cost of PF127 passivation versus mPEG/BSA passivation.

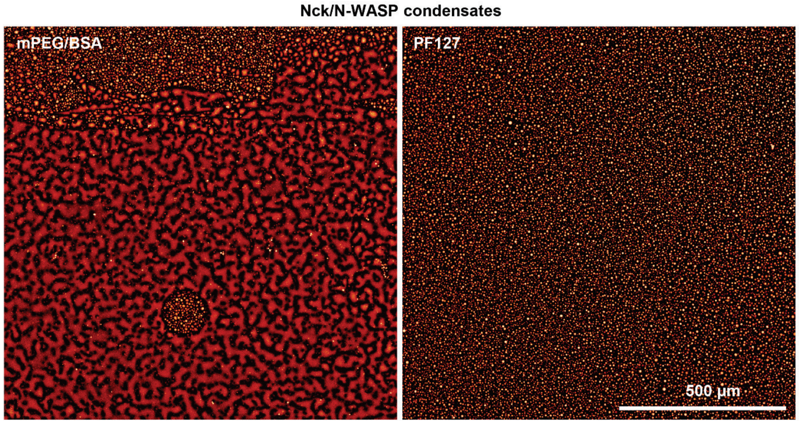


**Fig. S2. Characterization of homogeneity of PF127 passivation.**
Homogeneity comparison of PF127 and mPEG/BSA passivation in millimeter-scale fields of view.

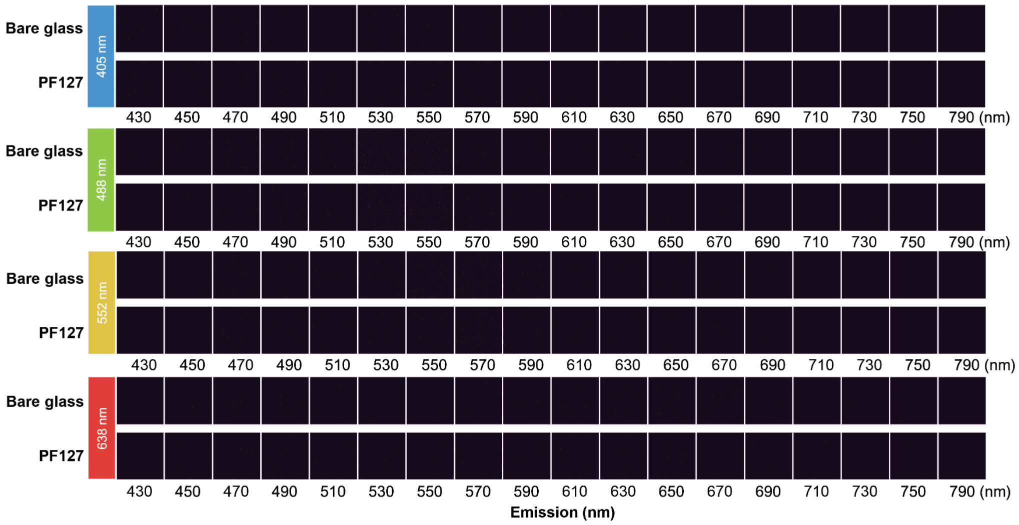


**Fig. S3. Homogeneity comparison of PF127 and mPEG/BSA passivation in millimeter-scale fields of view.**

Auto-fluorescence comparison of PF127-passivated glass with bare glass under 405 nm, 488 nm, 552 nm and 638 nm laser excitation. Emission light from 430 to 790 nm is monitored with a 20 nm bandwidth.

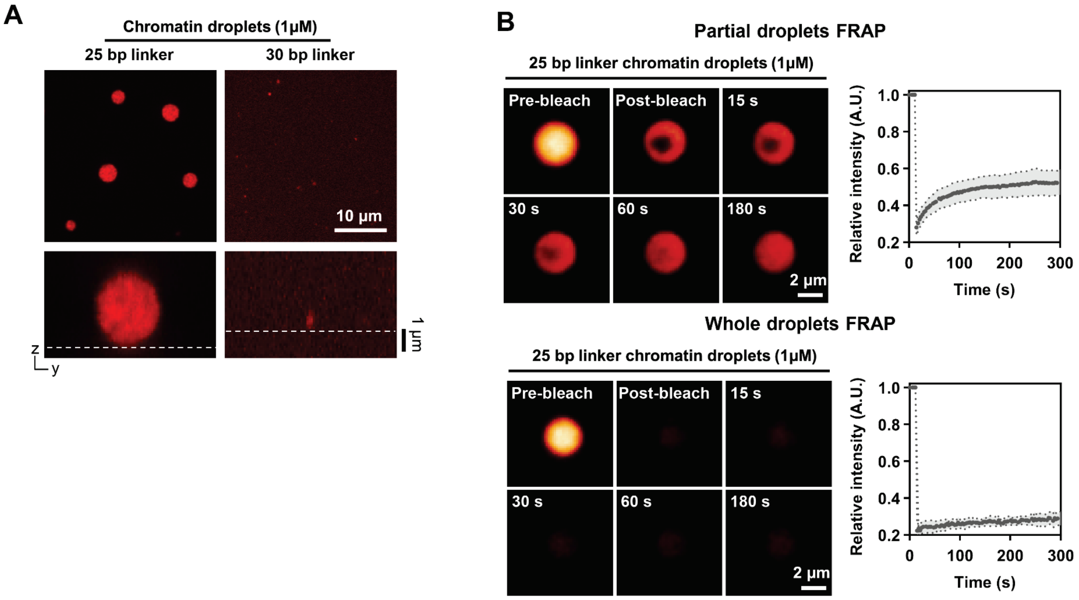


**Fig. S4. Characterization of phase separation behavior of chromatin condensates under PF127 passivation.**

A, Phase separation behavior of chromatin condensates with 25 bp and 30 bp linker lengths at 1 μM nucleosome concentration on PF127-passivated glass.

B, Partial and whole droplets FRAP of 25 bp linker chromatin condensates on PF127 passivated glass (mean ± SD, n=11 condensates for partial FRAP, n=5 condensates for whole droplets FRAP).

**Movie S1 (separate file).** Rotation and free movement of polySUMO/polySIM condensates across the PF127 passivated glass surface. Rectangular photo-bleached areas were used to observer rotation.

**Movie S2 (separate file).** Movement of polySUMO/polySIM condensates on PF127 passivated glass surface with different densities of anchor points, related to Fig. 4C.

**Movie S3 (separate file).** Trajectories of polySIM molecules within polySUMO/polySIM condensates, related to Fig. 5D. Images were captured at a 50 Hz rate. The playback speed is ten times the actual speed.

**Movie S4 (separate file).** Representative trajectories of mobile and immobile polySIM molecules within polySUMO/polySIM condensates, related to Fig. 5D.

**Movie S5 (separate file).** Trajectories of immobile polySIM molecules within polySUMO/polySIM condensates, related to Fig. 5E and 5F. Images were captured at a 50 Hz rate. The playback speed is ten times the actual speed.

**Movie S6 (separate file).** Trajectories of mobile polySIM molecules within polySUMO/polySIM condensates, related to Fig. 5E and 5F. Images were captured at a 50 Hz rate. The playback speed is ten times the actual speed.
